# Reducing food stress and parasitism may have unexpected consequences on faecal corticosterone in a wild rodent

**DOI:** 10.1101/2025.04.09.647990

**Authors:** S.E. Wolf, O. Dłużniewska, S.A. Babayan, A.B. Pedersen, T.J. Little

## Abstract

In wild animals, glucocorticoid hormones play a key role in responses to environmental stressors and the maintenance of homeostasis, allocating energy between energy-demanding processes. Although acute rises in glucocorticoids are beneficial for short-term survival, prolonged exposure can be detrimental to reproduction and survival. Thus, understanding the factors driving glucocorticoid concentrations is crucial for understanding key processes in ecology and evolution. Here, we examined how manipulation of two environmental stressors – food availability and parasite burden, influence levels of faecal corticosterone in a wild population of wood mice (*Apodemus sylvaticus*). To do so, we experimentally altered nutrition via food supplementation and reduced gastrointestinal nematode infection by anthelmintic treatment, using *Heligmosomoides polygyrus* as an indicator of treatment efficacy. Faecal corticosterone was not found to be impacted by food supplementation or anthelminthic treatment. However, our results may be mediated by variation in resource availability or masked by other factors that affect corticosterone. For example, corticosterone declined seasonally and was higher in adult, female mice. While we expected higher food availability and lower worm burden to decrease stress, putatively higher rates of reproduction in food supplemented areas could neutralise any decreases in corticosterone associated with alleviation of food and parasite stress. Thus, alleviating stress in the wild may have unintended consequences on fitness.

## 1. Introduction

Prolonged stress is detrimental to animal fitness (Defolie et al., 2020; Vitousek et al., 2019; Whirledge & Cidlowski, 2013). On a physiological level, the stress response is conveyed by glucocorticoids – steroid hormones released into circulation under the control of hypothalamic-pituitary-adrenal (HPA) axis (Timmermans et al., 2019). Working at the intersection of many physiological and behavioural processes, glucocorticoids are thought to regulate the allocation of energy resources to different functions, directing the trade-offs between key fitness related traits: growth, body maintenance, reproduction, and the immune response (Defolie *et al*., 2020; Landys *et al*, 2006; Patterson *et al*, 2014; Wingfield & Sapolsky, 2003).

Glucocorticoid release follows a circadian rhythm, helping maintain homeostasis according to diel and seasonal demands (Karaer *et al*., 2023; Spackman *et al*, 1978). Disruption of homeostasis by environmental stressors, however, can trigger elevation of glucocorticoid concentrations (Busch & Hayward, 2009). In response to short-term stress, glucocorticoids are thought to direct resources that promote survival, stimulating food-seeking behaviour and boosting the immune response (Cain & Cidlowski, 2017; Defolie *et al*., 2020; Dhabhar, 2014). In contrast, a chronic rise in glucocorticoids is associated with suppression of reproduction, growth, and immune function, the dysregulation of the latter making chronically stressed individuals more susceptible to infections (Defolie *et al*., 2020; Dhabhar, 2014; Edwards *et al*, 2019; Padgett & Glaser, 2003; Whirledge & Cidlowski, 2017; Wingfield & Sapolsky, 2003). While glucocorticoids are undoubtedly linked to fitness-related outcomes, how glucocorticoid levels are driven by environmental pressures warrants further study.

Glucocorticoid concentrations have also been associated with a variety of intrinsic factors. For example, sexual dimorphism in glucocorticoid actions appears partially driven by sex hormones (Handa et al., 1994; Kroon et al., 2020), in which oestrogen and testosterone stimulate and inhibit glucocorticoid secretion, respectively (Kalil et al., 2013). Along the same lines, glucocorticoids may become elevated during the breeding season due to the demands of reproduction (Romero, 2002), albeit potentially in a sex-specific manner (Putman et al., 2024). In addition, glucocorticoid responses to stress can differ by age (Heidinger et al., 2008; Wada et al., 2007; Wilcoxen et al., 2011), which may be driven by reproduction and/or senescence in adrenal capacity over a lifetime.

Infection by parasites (both macro- and microparasites/pathogens) is one very common environmental factor that can elicit a stress response via mechanisms such as behavioural changes, high energetic costs, and immune activation (Brace et al., 2017; Defolie et al., 2020; Wensveen et al., 2019). In fact, several meta-analyses show elevation of glucocorticoids in response to parasite infection in vertebrates (Defolie et al., 2020; O’Dwyer et al., 2020). However, the reverse relationship (St. Juliana et al., 2014; Sures et al., 2002) or a lack of change (Cizauskas et al., 2015; Monello et al., 2010; St. Juliana et al., 2014) have also been observed, suggesting that other factors such as host condition, coinfection, and the environment may mediate parasite-glucocorticoid relationships (Defolie et al., 2020; O’Dwyer et al., 2020).

Resource availability is another factor shown to influence glucocorticoids. Food restriction often results in higher glucocorticoid concentrations (Busch & Hayward, 2009; Feige-Diller et al., 2021; Heiderstadt et al., 2000; Kitaysky et al., 2001; Levay et al., 2010; Navarro-Castilla et al., 2019). For example, food limitation during El Niño conditions increased corticosterone levels in Galápagos marine iguanas, especially for individuals in poor condition (Romero & Wikelski, 2001). Short-term increases in glucocorticoids may be beneficial during periods of low resources by promoting gluconeogenesis (Sapolsky, 2000), the use of alternative energy sources like muscle protein, and food-seeking behaviours (Angelier et al., 2007; Kitaysky et al., 2001).

Such factors may also interact, for example, where the availability of resources may mediate how parasites affect glucocorticoid levels, or vice versa. Indeed, in the rodent *Peromyscus* spp., corticosterone reduction due to removal of intestinal parasites with anthelmintic treatment was significantly enhanced when food was supplemented (Pedersen & Greives, 2008). Higher food abundance and thus better nutritional status is linked to improved host immunity, faster recovery, and higher resistance to infections (Agostini *et al*, 2017; Becker *et al*, 2015; Blubaugh *et al*, 2023; Coop & Kyriazakis, 2001), while malnourished animals tend to have higher parasite burdens (Ezenwa, 2004). Still, only a limited number of studies examined the interaction between food availability, parasite burden, and glucocorticoids.

Here, we explored how variation in food availability and parasite burden predicted glucocorticoids in a wild population of wood mice, (*Apodemus sylvaticus*). Wood mice are commonly infected by a range of parasites, including the gastrointestinal (GI) nematode *Heligmosomoides polygyrus*, which is very prevalent (60 – 100%) (Gregory et al., 1990; Keymer & Dobson, 1987; Lewis et al., 2023). To identify the environmental drivers of glucocorticoids, we experimentally (1) increased high quality food availability and (2) reduced *H. polygyrus* burdens through anthelmintic treatment. We then measured faecal corticosterone, which is known to reflect blood hormone levels over a period of hours to days (Frynta *et al*, 2009; Karaer *et al*., 2023; Palme, 2019). We predicted that increased food availability and decreased parasite burden would alleviate stress, potentially in an additive manner, and lower faecal corticosterone concentrations.

## 2. Materials And Methods

### 2.1 Ethics Statement

All animal work was carried out under the approved UK Home Office Project License PP4913586 in accordance with the UK Home Office in compliance with the Animals (Scientific Procedures) Act 1986 and approved by the University of Edinburgh Ethical Review Committee. Fieldwork was performed with permission from landowners.

### 2.2 Field Work, Mouse Capture, and Sample Collection

Wild wood mice were captured between 30 May 2023 and 23 November 2023 at two forested sites near Edinburgh, Scotland, United Kingdom: Penicuik Estates (55.8261° N, 3.2467° W) and Bilston Wood (55.8714° N, 3.1470° W). At each site, two 80m x 80m grids were placed at least 50m apart. Each grid was divided into 64 trap stations (10m x 10m), each containing two Sherman live traps (H.B. Sherman 2 × 2.5 × 6.5 in. folding traps, Tallahassee, FL, USA), set under trap covers to protect animals from inclement weather; resulting in 256 traps set each trapping night.

We live-trapped each woodland site every three weeks for three consecutive nights for a total of 9 trapping sessions (defined as 3 consecutive trapping nights in a week). Sherman live traps were set late each afternoon and collected the following morning. Each trap contained bedding and a bait consisting of a mix of seeds, mealworms, and a carrot slice. At first capture, individuals above 12g were given subcutaneous RFID tags (125kHz, Francis Scientific Instruments, UK). Individuals smaller than 12g were given a unique ear biopsy, and when recaptured at a later date and they were >12g, received a microchip.

Upon first and all subsequent captures, several individual traits were recorded and then wood mice were released at site of capture. Sex was determined based on the appearance of genitalia. To determine reproductive condition, for males, individuals with descended and scrotal testes were considered reproductive, while males with abdominal (non-visible) testes were non-reproductive. Lactating or pregnant females were reproductive, while females with perforate or non-perforate vagina were non-reproductive (i.e., clearly not pregnant or rearing young). Approximate age (juvenile, subadult, adult) was determined based on body mass (juvenile: <10g; subadult: 10-15g, adult >15g) and fur colour, which transitions from grey to brown pelage into adulthood.

Faecal samples were collected from clean, autoclaved Sherman live traps within 24-48 hours of capture. First, faecal pellets were weighed and stored in formalin (4% formaldehyde) at 4°C until faecal eggs counts were performed. We also collected 5-10 dry faecal pellets in an Eppendorf tube (when available), which was frozen immediately at -80°C until faecal steroid extraction.

### 2.3 Food Supplementation and Drug Treatment

We used a two-way factorial experimental design to manipulate both food availability and GI nematode burdens. First, at each of two sites, one grid was food-supplemented and one was given no additional food; mice on both grids had access to their normal diet. On food supplemented grids, we distributed 10kg of TransBreed^®^ or Safe^®^ 199-high quality nutrient veterinary feeds (protein 18.5%, fat 9.5%, starch 36.8%, high micronutrients; see Sweeny et al., 2021) evenly over the grid once a week, starting two weeks before the first trapping session. Parasite treatment was conducted at the individual level. Wood mice captured on all 4 grids were randomly (but balanced within each sex) assigned at first capture to receive either anthelminthic drug treatment or a water control. Individuals assigned drug treatment received a combination of 100kg/mg Pyrantel and 9.4kg/mg Ivermectin using a sterilised oral gavage. Ivermectin and Pyrantel are common anthelmintic drugs that reduce gastrointestinal nematodes (Pedersen & Fenton, 2015); this combination of drug treatments reduces *H. polygyrus* burdens by >99% in wild wood mice for ∼16 days (Clerc et al., 2019; Sweeny et al., 2021). Due to delayed delivery of Pyrantel, drug treatment only included the single drug dose of 9.4 kg/mg Ivermectin; however, after this point all drug treated individuals received the combination of drugs. This dose of Ivermectin alone can reduce the probability of nematode infection by >70% (Knowles et al., 2012); therefore, we are confident that this small group of individuals experienced significant decreases in *H. polygyrus* after this first dose. Only individuals larger than 12g were RFID-tagged and received treatment, and both anthelmintic and control treatments were repeatedly given at all subsequent recaptures, with a minimum time between doses of 21 days.

### 2.4 Faecal Egg Counts

Faecal samples were analysed using a modified cuvette method (adapted from Hayward et al., 2019). First, the samples were homogenised and then spun down (1500 rpm for 2 minutes) in thin-walled polypropylene tubes, from which the supernatant was aspirated off. The remaining compact faecal matter was thoroughly mixed with a saturated salt solution and spun down again (1500 rpm for 2 minutes). The thin-walled tubes were clamped just below the meniscus using medical forceps, and this top layer of liquid containing GI parasite eggs was poured into a fecalizer. We allowed oocysts to float onto a coverslip placed on top of the fecalizer for 20 minutes, at which time it was placed onto a slide for analysis. *H. polygyrus e*ggs were counted at 10x magnification. Egg counts were adjusted by the weight of the faecal sample to generate an ‘eggs per gram’ of faeces (EPG). EPG were quantified for all available faecal samples from all animals/captures (n = 888, n = 1 – 20 samples per individual).

### 2.5 Faecal Steroid Extraction

Faecal corticosterone was extracted from frozen faecal samples from all resident individuals, i.e., those sampled for 2 or more trapping sessions. This resulted in 268 total samples from 100 individuals; with 1 to 8 faecal corticosterone samples from each individual over the course of the field season. The faecal steroid extraction protocol was adapted from (Nomoto & Kansaku, 2023; Veitch *et al*., 2021). Frozen faecal samples (4-5 pellets) were thawed and dried in a heating block at 70°C and powdered using a pestle homogenizer. The extraction was performed by adding 1 ml of freshly prepared 80% (v/v) methanol solution to approx. 0.005 g of faecal powder. Fully submerged samples were vortexed for 10s and shaken overnight at room temperature on a tube shaker. Samples were then centrifuged for 10 min at 10,000*g*. 0.8mL of supernatant were transferred to a new tube and the liquid was evaporated in a heating block at 70°C to isolate steroids. Extracted steroids were stored at -20°C until the assay, when they were reconstituted in [400 - 800 µL] immunoassay buffer (Abcam). Samples were further diluted if necessary.

### 2.6 Corticosterone ELISA

Faecal corticosterone concentrations were quantified through a competitive enzyme-linked immunosorbent assay (ELISA) using the Abcam Corticosterone ELISA kit (AB108821) following the manufacturer’s instructions. Absorbance was read at 450 nm with the wavelength correction at 570 nm using FLUOstar Omega microplate reader (BMG Labtech). Corticosterone concentrations were extrapolated from the standard curve generated with a 4-Parameter fit using Omega Data Analysis software (version 3.32 R5). Corticosterone concentrations were standardised against the weight of faecal samples and presented as ng of faecal corticosterone per g faeces (ng/g). Samples with a high coefficient of variation (CV >20%) were rerun. The failure rate was 7.6% with samples being excluded due to a high CV or falling below the standard curve range.

Kit sensitivity was 0.26 ng/ml. Kit cross reactivity with other steroids was as follows: progesterone <2%, allopregnanolone 1%, cortisone <0.5%, aldosterone (<0.3%), 4-Pregnen-17, 20β-Diol-3-One (<0.12%), and below 0.1% for other steroids, including testosterone and cortisol.

The average intra-plate variation was 8.9% and the inter-plate variation was 39.8%. However, we are confident that our results are not driven by this inter-plate variation. First, faecal samples were assayed in duplicate with all samples from the same individual tested on the same plate, unless singular samples had to be re-measured. All plates, except for plates containing sample reruns, were also balanced in terms of food treatment, drug treatment, sex, age, body mass, and date. Second, we accounted for inter-plate variation in all models by adding plate ID as a random effect. Third, we reran all models using plate-adjusted corticosterone values, in which each sample value was divided by the corticosterone value of its inter-plate control sample and then multiplied by the plate-wide average corticosterone value. This analysis did not significantly change model results (see **Supplementary Material Tables S1, S3, and S5**).

### 2.7 Statistical Analyses

Statistical analyses were done in R studio using the ‘lme4’ package for (general) linear mixed-effects models (Bates et al., 2014).

We first assessed the effects of food supplementation and anthelminthic treatment on several metrics related to reproduction and body condition. First, abundance of *Apodemus* was measured as the **minimum number known alive** (MNKA) for each of the four treatment combinations (supplementation x anthelminthic treatment) during trapping sessions denoted as integers 1 – 9 (e.g., May to November 2023, following Wolff 1996). To test for the effects of food supplementation and anthelminthic treatment on *Apodemus* abundance, we used a glm to predict MNKA per treatment combination and trapping session, with fixed effects of food and anthelminthic treatment, and their interaction, as well as trapping session. Second, we tested effects on **reproductive status**. We used a binomial general linear mixed effect model, with reproductive status as the response variable and fixed effects of food and anthelminthic treatments, as well as their interaction, sampling date, sex, age, and trapping location, with ID as a random effect. Last, we tested for effects on two proxies of condition: **residual body index** (i.e., the difference between an individual’s actual and predicted mass:length ratio) and **skeletal muscle index** (i.e., an estimate of muscle mass). For each variable, we ran a linear mixed effect model with fixed effects of food and worm egg burden, as well as their interaction, anthelminthic treatment, trapping date, sex, age, reproductive status, and trapping location, with ID as a random effect.

Next, *H. polygyrus* egg burden (eggs/gram faeces) was log(x+1)-transformed and rounded to the nearest integer. Using this egg burden as count data, we ran a general linear-mixed effect model (*glmer*) assuming a Poisson distribution. We included food supplementation (no supplementation, supplementation) and egg burden as fixed effects. In addition, we also included anthelminthic treatment (control, anthelminthic) as a fixed effect to account for additional effects of the drug (e.g., alterations in gut community). Other fixed effects included sampling date, reproductive status (reproductive, non-reproductive), sex (male, female), and age category (non-adult, adult). Both juveniles and subadults were considered non-adults. Sampling date was formatted as Julian date, i.e., days since 1 Jan, and was scaled from 0 to 1 during modelling to reach model convergence. Additionally, to account for variation between two spatial replicates, we included site as a fixed effect. We also included a random effect of mouse ID to account for repeated samples from the same individual.

To address how manipulations of food availability and parasite burdens influence levels of corticosterone, we tested for the additive effect of anthelmintic treatment and food supplementation on faecal corticosterone levels (ng/g), which were log-transformed. We ran a linear mixed effects model assuming a gaussian distribution that included the same fixed effects as those in the previous (egg burden) model: food supplementation, anthelminthic treatment, and their interaction, as well as sampling date, reproductive status, sex, age category, and site. Random effects of mouse ID and ELISA plate ID accounted for repeated samples from the same individual and inter-plate variation, respectively. Residuals of this model were approximately normal. Test statistics were calculated using Type II sum of squares and reported with Kenward-Roger degrees of freedom. Similar linear mixed effects models were run to explore further variation in anthelminthic drug treatment: (1) a model which included only faecal samples collected post-treatment (i.e., no first captures, as the drug treatment was given on the first capture and therefore its effects on egg burdens would not be detectable until subsequent captures) and one model that replaced anthelminthic treatment with the number of doses received up to the sampling date, to test for a cumulative effect of multiple drug doses (**See Supplementary Material Tables S2 – S5**).

## 3. Results

From May to November 2023, 317 unique wood mice were captured across our 4 experimental grids, 286 of which were adults and were treated with either water or anthelminthic (Ant) drugs (Group sizes: Food-Ant = 84; Food-Con = 83; Con-Ant = 59; Con-Con = 60). We found that food supplemented grids had a higher abundance of wood mice compared to non-supplemented grids (X^2^ = 25.50, p = < 0.0001, **Fig 1A**), as well as a higher number of juvenile mice captured across our grids, although this was not significant (X^2^ = 18.00, p = 0.051, **Fig 1A**). While neither food supplementation or anthelmintic treatment was associated with the number of reproductive individuals on our grids (supplementation: X^2^ = 0.47, p = 0.49; anthelminthic: X^2^ = 0.06, p = 0.80), mice in reproductive condition were significantly more likely to be found earlier in the season (X^2^ = 66.69, p < 0.0001), and were more likely to be male (X^2^ = 49.14, p < 0.0001) and adult (X^2^ = 36.55, p < 0.0001). In addition, our treatments did not have an effect on residual body index or skeletal muscle index, although female, reproductive, adult individuals tended to have higher body condition-related indices (**See Supplemental Materials Tables S6, S7**).

**Figure 1.**
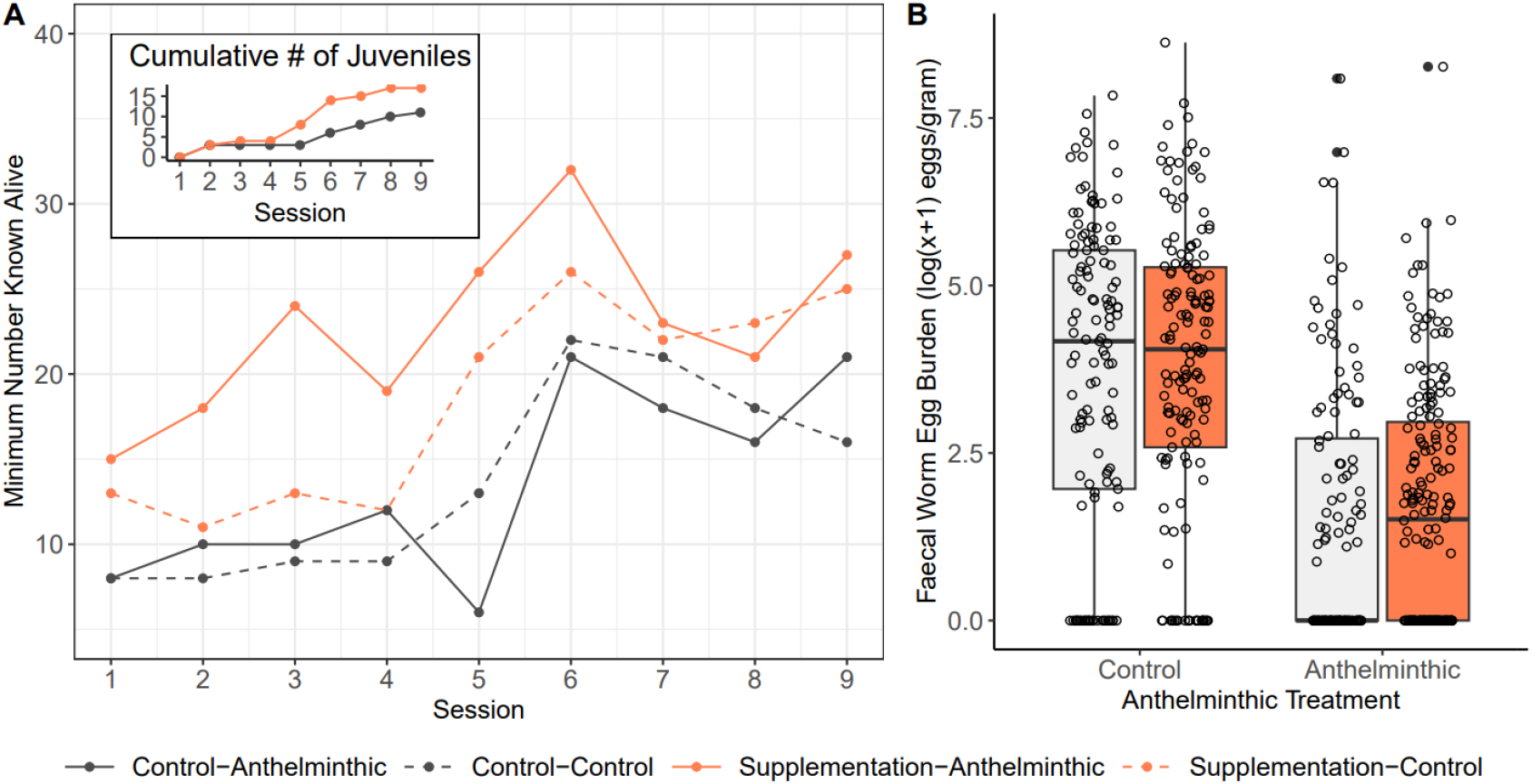
Main treatment effects of food supplementation and anthelminthic treatment. (A) Population dynamics: wood mouse abundance measured as the minimum number known alive per each of four treatment combinations from May to November 2023. The inset shows the cumulative number of juveniles found on control (black; n = 2 grid replicates) and food supplemented (orange; n = 2 grid replicates) grids from May to November 2023. (B) The effect of food and anthelminthic treatments on *H. polygyrus* egg burden in wood mice, measured and plotted as raw log(x+1)-transformed eggs per gram faeces, and points represent individual samples.

We assessed 888 faecal samples for *H. polygyrus* egg burden, which ranged from 0 to 5578 eggs per gram faeces (EPG) (mean = 123.69 ± 12.35 EPG; median = 13.33 EPG). Anthelminthic treatment significantly reduced average *H. polygyrus* egg burdens by 52% (**Table 1, Fig 2B**), whereas control individuals increased in average egg burden by 80% from first to subsequent captures later in the season. Similarly, anthelminthic-treated mice exhibited a 17% decrease in *H. polygyrus* prevalence (first capture = 67%, subsequent captures = 55%), whereas control prevalence increased by 23.35% (first capture = 66%, subsequent captures = 82%). *H. polygyrus* EPG was however, not impacted by food supplementation, sampling date, reproduction, sex, or age.

**Table 1.** The effect of food supplementation and anthelminthic treatment on *H. polygyrus* egg burdens in wood mice. Eggs per gram (EPG) is log(x+1)-transformed and rounded to the nearest integer before being run in a general linear mixed model with a Poisson distribution. Mouse ID is included as a random effect. Reference levels include: no food supplementation, no anthelminthic treatment, and non-reproductive, male, and non-adult individuals. ^#^p < 0.1, *p < 0.05.

| Fixed Effects | Estimate ± SE | z-value | p-value |
| --- | --- | --- | --- |
| Intercept | 0.21 ± 0.39 | 0.54 | 0.59 |
| Food Availability | 0.13 ± 0.16 | 0.79 | 0.43 |
| Drug Treatment | -0.41 ± 0.18 | -2.21 | 0.03* |
| Julian Date<sup>1</sup> | 0.60 ± 0.39 | 1.53 | 0.13 |
| Reproductive Status | 0.03 ± 0.11 | 2.78 | 0.78 |
| Sex | 0.008 ± 0.13 | 0.06 | 0.95 |
| Age | 0.14 ± 0.15 | 0.94 | 0.35 |
| Site | 0.32 ± 0.12 | 2.53 | 0.01* |
| Food x Egg Burden | -0.14 ± 0.23 | -0.63 | 0.53 |

<sup>1</sup> Julian Date is scaled from 0 (end of May) to 1 (end of November)

**Figure 2.**
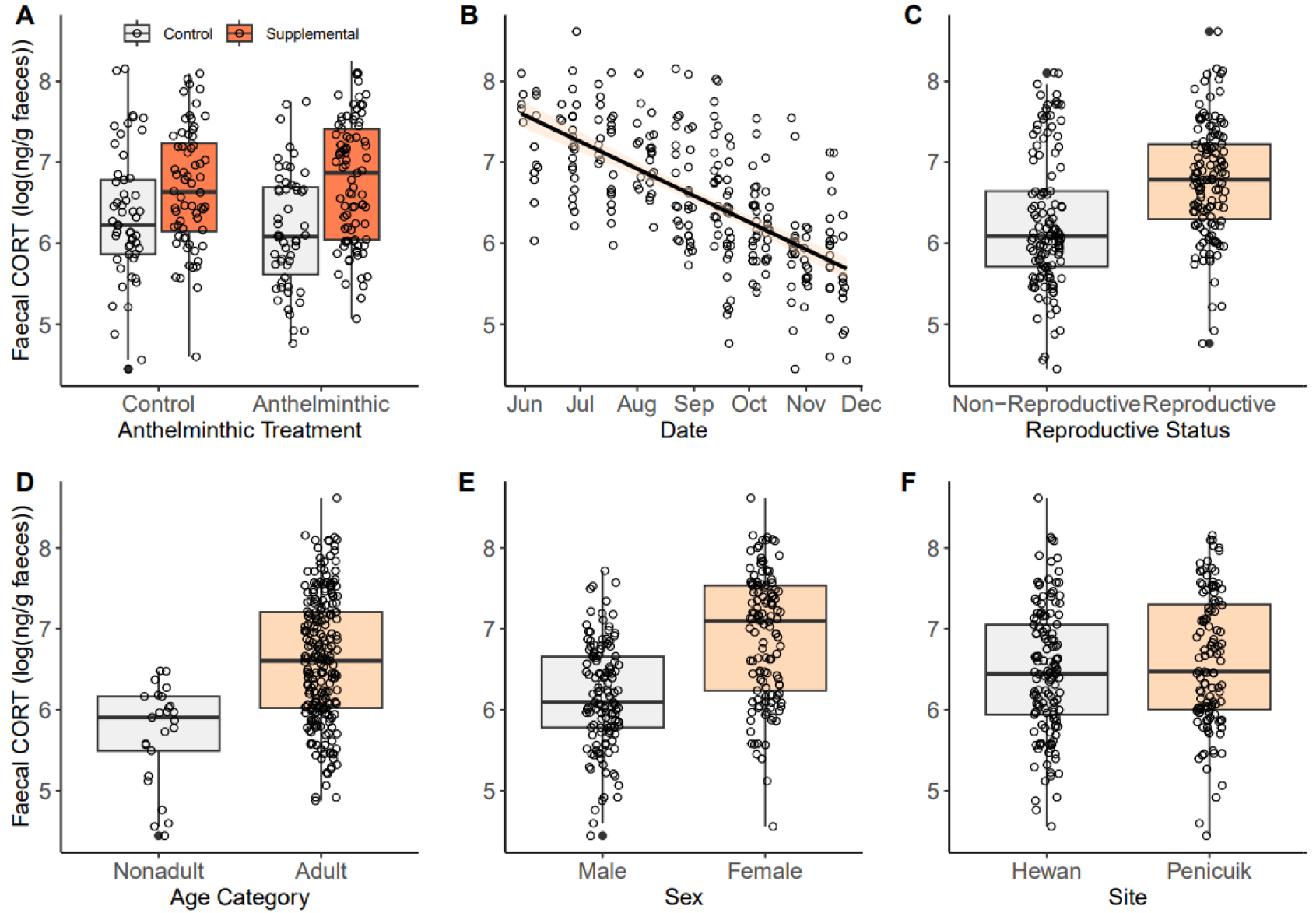
Predictors of faecal corticosterone in a wild population of wood mice. Predictors include: (A) experimental manipulation of food availability (high quality mouse chow vs no supplementation) and anthelmintic treatment to reduce *H. polygyrus* burdens (treatment with Ivermectin and Pyrantel vs water control); (B) date of faecal sample collection; (C) reproductive status; (D) age category; (E) sex; and (F) location of sampling. Plots are based on data from 268 samples and 100 individuals, and points represent individual samples. Faecal corticosterone concentration is plotted as predicted values from the linear mixed-effects model based on the log-transformed concentration (ng/g).

We successfully assessed 268 samples for faecal corticosterone from 100 mice (52 male, 48 female; captured in 2^+^ trapping sessions across the field season; therefore considered residents). Faecal corticosterone levels ranged from 35.57 to 23,406.06 ng/g (mean = 1724.98 ± 190.89 ng/g; median = 616.91 ng/g). While faecal corticosterone levels were somewhat higher in wood mice on food supplemented grids (p = 0.086, mean_control_ = 6.26, mean_food_ = 6.71 log(ng/g), we found no evidence for a significant effect of either food supplementation or anthelminthic treatment (**Table 2, Fig 2A**).

**Table 2.** Linear mixed-effects model of the effect of food supplementation and anthelminthic treatment on faecal corticosterone (log-transformed ng/g) in wood mice. Data is based on 268 samples from 100 individuals, and mouse ID and assay plate ID were included as random effects. Test statistics are calculated using Type II sum of squares and reported with Kenward-Roger degrees of freedom. Reference levels include: no food supplementation, no anthelminthic treatment, and non-reproductive, male, and non-adult individuals. ^#^p < 0.1, *p < 0.05.

| Fixed Effects | Estimate ± SE | df | F-value | p-value |
| --- | --- | --- | --- | --- |
| Intercept | 7.71 ± 0.56 |  |  |  |
| Food Availability | -0.01 ± 0.23 | 1, 100.20 | 2.99 | 0.086<sup>#</sup> |
| *H. polygyrus* Egg Burden | -0.09 ± 0.05 | 1, 236.65 | 1.38 | 0.24 |
| Drug Treatment | -0.10 ± 0.16 | 1, 75.00 | 0.37 | 0.54 |
| Julian Date<sup>1</sup> | -2.94 ± 0.53 | 1, 212.31 | 29.74 | <0.0001* |
| Reproductive Status | 0.33 ± 0.17 | 1, 213.37 | 3.79 | 0.052<sup>#</sup> |
| Sex | 0.58 ± 0.16 | 1, 83.40 | 12.08 | 0.001* |
| Age | 0.73 ± 0.25 | 1, 210.97 | 8.37 | 0.004* |
| Site | 0.32 ± 0.16 | 1, 72.27 | 4.05 | 0.04* |
| Food x Egg Burden | 0.10 ± 0.06 | 1, 233.45 | 2.57 | 0.11 |

<sup>1</sup> Julian Date is scaled from 0 (end of May) to 1 (end of November)

Additionally, no effect of anthelminthic treatment was found when subsetting the data to include only faecal samples collected post-drug treatment (i.e., only including post-treatment captures) or when controlling for the number of doses received up to the sampling date (**See Supplementary Tables S2 – S5**). However, the date of sampling significantly predicted faecal corticosterone (**Fig 2B**), which decreased in concentration from June to November. Reproductive mice had higher, but non-significant levels of corticosterone than non-reproductive mice (p = 0.052, mean_non_ = 6.27, mean_reproductive_ = 6.78 log(ng/g) (**Fig 2C**). We also found that faecal corticosterone was higher in female and adult mice (**Fig 2D, 2E**). Lastly, there was no effect of the woodland site (**Fig 2F**).

## 4. Discussion

Identifying factors driving glucocorticoid levels may improve our understanding of the causes and consequences of variation in survival and fitness. Resource availability and immune challenges and or pathology due to parasite infection can be considerable stressors in wild animals that may alter glucocorticoid levels in an additive manner. However, we show no effects of food supplementation, anthelminthic treatment, or their interaction on faecal corticosterone levels in wild wood mice (*Apodemus sylvaticus*). While previous work has shown a robust effect of these manipulations on corticosterone in *Peromyscus* spp. (Pedersen & Greives, 2008), effects here may be mediated by year-to-year variation in baseline resource availability (e.g., masting cycles of trees) or masked by other drivers of corticosterone like seasonal changes, sex, reproductive activity, and age. In addition, food supplementation and anthelminthic treatment may result in antagonistic effects on corticosterone that produce an overall null result in this system. For example, increased reproduction is one putative byproduct of higher food availability that could contribute to elevated corticosterone and effectively neutralise any decreases in corticosterone associated with alleviation of food and parasite stress. Below, we start to explore these complex interactions between environmental and intrinsic factors that may be important contributors to variation in fitness in the wild.

Our results contrast with observations from *Peromyscus* spp., in which anthelminthic drug treatment alone and with food supplementation significantly lowered glucocorticoid levels (Pedersen & Greives, 2008). However, lack of corticosterone response to parasite removal has been observed in other studies (Carlsson et al., 2016; Monello et al., 2010; Trevisan et al., 2017), suggesting that some macroparasites like *H. polygyrus* may be tolerated by the host during chronic infections, thereby putting less pressure on the immune system and resulting in an acute rise in glucocorticoids (Defolie *et al*., 2020; Monello *et al*., 2010; O’Dwyer *et al*., 2020). Anthelmintic-driven removal of a prevalence gastrointestinal nematode has also been shown to increase the abundance of other co-infecting pathogens (Pedersen & Antonovics, 2013), implying that the relative infection stress in our study may have remained unchanged even after *H. polygyrus* was reduced. Alternatively, in our current study only individuals captured 2+ times were used for corticosterone analyses. Although this was necessary to examine the effects of drug treatment, it may have inadvertently biased results towards individuals in better condition that survived to be captured on multiple occasions, something to be explored further.

Additionally, the time of our study may have contributed to the observed relationships or lack thereof. In *Peromyscus* spp., corticosterone response to anthelmintics and provisioning was only detectable outside the reproductive season (Pedersen & Greives, 2008). Our samples were collected predominantly during late summer, when wood mice commonly reproduce (Kufner & Moreno, 1988). Wood mice, as small and short-lived animals, may prioritise reproduction over the immune response and other energy-demanding processes, as maximising reproductive output benefits their overall fitness (Díaz & Alonso, 2003). Indeed, food supplementation has been demonstrated to increase reproductive activity at a cost to body condition (Díaz & Alonso, 2003). Reproducing individuals often carry higher parasite burdens (Shaner *et al*, 2018), further supporting trade-offs between reproduction and immunity, which may be particularly visible when resources are scarce (Viney & Riley, 2017). In our study, food-supplemented grids had more mice, which may suggest that reproductive activity was higher. That there were not more reproductive individuals on those grids may reflect limitations in our ability to identify females during early stages of pregnancy. Regardless, if additional resources were directed towards reproduction, our observed corticosterone levels on food supplemented grids may reflect increased energy demands caused by reproductive activity. Similarly, reproduction may partially contribute to higher corticosterone in adult mice versus nonadults, which are less likely to reproduce.

Despite the fact that high glucocorticoid concentrations can suppress reproduction (Whirledge & Cidlowski, 2017), reproduction itself is costly and can act as a stressor (Michael Romero, 2002). Although reproductive status was not significant in our model, possibly due to overlapping variation with age and sex, faecal corticosterone appears higher in reproducing than non-reproducing individuals. Previous work suggests that sex differences in corticosterone may result from the effects of male and female sex hormones on the HPA axis during the breeding season. Female oestrogens stimulate the HPA axis and corticosterone release, while male testosterone acts as an inhibitor (Handa *et al*, 1994). Furthermore, as glucocorticoids can be especially high in pregnant or lactating females (Karaer *et al*., 2023; Palme, 2019), this direct cost of reproduction may make females more dependent on resources and have higher energy demands and food intake, which could elevate corticosterone (Navarro-Castilla et al., 2019).

Corticosterone levels decreased over the course of the study from June - November. Seasonal glucocorticoid fluctuations are observed in many species (Dantzer *et al*, 2016; David *et al*, 2015; Lanctot *et al*, 2003; Puehringer-Sturmayr *et al*, 2018; Zhou *et al*., 2020). Although this may reflect changes in weather conditions, food abundance, or parasite prevalence (Michael Romero, 2002), corticosterone concentrations typically maximise during the breeding season, suggesting that the reproduction cost may drive seasonal changes in glucocorticoids. In our study, corticosterone concentrations were the highest during the summer and started to drop in September, which was reflected by a similar decrease in reproductive individuals. Since similar seasonal patterns of glucocorticoids have previously been observed in wood mice (Sánchez-González *et al*, 2018), this may suggest that reproductive activity contributes to seasonal corticosterone variation observed in this study.

Clearly, further research will be necessary to get a full picture of the interactions between infection burden, immune response, reproduction, and resource abundance. This may include looking directly at the parasite – corticosterone relationship as well as evaluating sex hormones and immune parameters as potential downstream effectors of corticosterone. With glucocorticoids having prominent and pleiotropic effects, investigation of the drivers of their natural variation may be crucial for understanding population dynamics and drivers of variation in fitness.

## Acknowledgements

We are grateful to Mairéad Corr, Alexandra Vavrik, Thomas Loebel-Messer, Isabel Entwistle, Rhoslyn Howroyd, Anders Erlandson, Cara Duffy, and Nathan Loebel-Messer for support in the field and laboratory; to Cameron Watkins and James Nixon for access to field sites; and to our reviewers for feedback. This research was supported by a NERC grant (NE/X001423/1) to TL, ABP, and SAB and the Wellcome Trust (Host, Pathogens and Global Health PhD programme; 218492/Z/19/Z) studentship to OD.

## Declaration of Competing Interest

The authors declare no conflict of interest.

## Data Availability Statement

Data will be available on the Dryad Digital Repository.

## Author Contributions

**Sarah E. Wolf**: Conceptualization, Data curation, Formal Analysis, Investigation, Methodology, Project administration, Supervision, Visualization, Writing – original draft, Writing – review and editing. **Olga Dłużniewska**: Data curation, Formal Analysis, Investigation, Methodology, Visualization, Writing – original draft, Writing – review and editing. **Simon A. Babayan**: Conceptualization, Funding acquisition, Supervision, Writing – review and editing. **Amy B. Pedersen**: Conceptualization, Funding acquisition, Resources, Supervision, Writing – review and editing. **Tom J Little**: Conceptualization, Funding acquisition, Investigation, Resources, Supervision, Writing – review and editing.

## Notes

### Competing Interest Statement

The authors have declared no competing interest.

