## Supplement for "Reducing food stress and parasitism may have unexpected consequences on faecal corticosterone in a wild rodent"

**Tables S1 – S5:** Linear mixed-effects models of the effect of food supplementation, anthelminthic treatment, and other factors on faecal corticosterone (log-transformed ng/g) in wood mice. For all models, mouse ID and assay plate ID were included as random effects. Test statistics are calculated using Type II sum of squares and reported with Kenward-Roger degrees of freedom. Reference levels: no food supplementation, no anthelminthic treatment, and non-reproductive, male, and non-adult individuals.

^#^p < 0.1, *p < 0.05.

Models vary in two main ways:

1. Tables S2 – S5 aim to further explore the predictive power of anthelminthic treatment. Tables S2 – S3 exclude samples taken during first captures, at which time individuals have not received anthelminthic or control treatments. Tables S4 – S5 replace anthelminthic treatment with the number of times each individual was treated before sample collection.
2. Tables S1, S3, and S5 aim to adjust for inter-plate variation in corticosterone values. Here, each sample value was divided by the corticosterone value of its inter-plate control sample and then multiplied by the plate-wide average corticosterone value. This brings all corticosterone values towards the study-wide average and minimizes inter-plate variation.

Readers should note that anthelminthic treatment never significantly predicts corticosterone values, regardless of type of anthelminthic variable / subset and regardless of adjustments for inter-plate variation in corticosterone values.

| **Table S1:** All corticosterone values, corticosterone values adjusted for inter-plate variation (n = 268 samples from 100 individuals). | | | | |
| --- | --- | --- | --- | --- |
| **Fixed Effects** | **Estimate ± SE** | **df** | **F-value** | **p-value** |
| Intercept | 8.55 ± 0.56 |  |  |  |
| Food Availability | 0.02 ± 0.23 | 1, 101.49 | 3.10 | 0.08^#^ |
| Egg Burden | -0.09 ± 0.05 | 1, 238.04 | 1.44 | 0.23 |
| Drug Treatment | -0.11 ± 0.16 | 1, 75.90 | 0.49 | 0.48 |
| Julian Date^1^ | -2.96 ± 0.53 | 1, 214.15 | 30.09 | <0.0001* |
| Reproductive Status | 0.32 ± 0.17 | 1, 215.32 | 3.51 | 0.06^#^ |
| Sex | 0.56 ± 0.16 | 1, 84.06 | 11.50 | 0.001* |
| Age | 0.81 ± 0.25 | 1, 212.04 | 10.06 | 0.002* |
| Site | 0.31 ± 0.16 | 1, 72.96 | 4.00 | 0.049* |
| Food x Worm Burden | 0.09 ± 0.06 | 1, 234.47 | 2.08 | 0.15 |
| ^1^Julian Date is scaled from 0 (end of May) to 1 (end of November) | | | | |
| **Table S2:** Subsetted to exclude pre-treatment (first capture) samples (n = 175 samples from 93 individuals). | | | | |
| **Fixed Effects** | **Estimate ± SE** | **df** | **F-value** | **p-value** |
| Intercept | 7.95 ± 0.76 |  |  |  |
| Food Availability | 0.16 ± 0.29 | 1, 75.74 | 3.73 | 0.057^#^ |
| Egg Burden | -0.08 ± 0.06 | 1, 157.30 | 0.75 | 0.39 |
| Drug Treatment | -0.12 ± 0.19 | 1, 57.53 | 0.40 | 0.53 |
| Julian Date^1^ | -3.34 ± 0.70 | 1, 151.38 | 22.33 | <0.0001* |
| Reproductive Status | 0.48 ± 0.20 | 1, 142.76 | 5.48 | 0.02* |
| Sex | 0.73 ± 0.20 | 1, 67.70 | 13.29 | 0.001* |
| Age | 0.72 ± 0.37 | 1, 148.44 | 3.68 | 0.057^#^ |
| Site | 0.10 ± 0.18 | 1, 57.69 | 0.29 | 0.59 |
| Food x Worm Burden | 0.07 ± 0.08 | 1, 150.66 | 0.78 | 0.38 |
| ^1^Julian Date is scaled from 0 (end of May) to 1 (end of November) | | | | |

| **Table S3:** Subsetted to exclude pre-treatment (first capture) samples and corticosterone values adjusted for inter-plate variation (n = 175 samples from 93 individuals). | | | | |
| --- | --- | --- | --- | --- |
| **Fixed Effects** | **Estimate ± SE** | **df** | **F-value** | **p-value** |
| Intercept | 8.77 ± 0.75 |  |  |  |
| Food Availability | 0.20 ± 0.29 | 1, 77.38 | 3.60 | 0.06^#^ |
| Egg Burden | -0.07 ± 0.06 | 1, 158.68 | 0.81 | 0.37 |
| Drug Treatment | -0.13 ± 0.19 | 1, 58.82 | 0.48 | 0.49 |
| Julian Date^1^ | -3.36 ± 0.69 | 1, 153.92 | 22.76 | <0.0001* |
| Reproductive Status | 0.46 ± 0.20 | 1, 145.76 | 5.06 | 0.03* |
| Sex | 0.72 ± 0.19 | 1, 68.81 | 13.11 | 0.001* |
| Age | 0.81 ± 0.37 | 1, 149.98 | 4.67 | 0.03* |
| Site | 0.10 ± 0.18 | 1, 57.92 | 0.27 | 0.60 |
| Food x Worm Burden | 0.06 ± 0.08 | 1, 153.24 | 0.44 | 0.51 |
| ^1^Julian Date is scaled from 0 (end of May) to 1 (end of November) | | | | |

| **Table S4:** All corticosterone values, replacing anthelminthic treatment with the number of treatments received prior to each sample (n = 268 samples from 100 individuals). | | | | |
| --- | --- | --- | --- | --- |
| **Fixed Effects** | **Estimate ± SE** | **df** | **F-value** | **p-value** |
| Intercept | 7.67 ± 0.58 |  |  |  |
| Food Availability | -0.04 ± 0.23 | 1, 99.82 | 2.77 | 0.09^#^ |
| Egg Burden | -0.09 ± 0.05 | 1, 238.46 | 1.03 | 0.31 |
| Number of Treatments Received | 0.006 ± 0.06 | 1, 100.41 | 0.01 | 0.92 |
| Julian Date^1^ | -2.95 ± 0.59 | 1, 169.76 | 24.58 | <0.0001* |
| Reproductive Status | 0.33 ± 0.17 | 1, 220.45 | 3.49 | 0.059^#^ |
| Sex | 0.58 ± 0.16 | 1, 83.16 | 12.10 | 0.001* |
| Age | 0.72 ± 0.25 | 1, 209.74 | 7.96 | 0.01* |
| Site | 0.31 ± 0.16 | 1, 72.65 | 3.83 | 0.054^#^ |
| Food x Worm Burden | 0.11 ± 0.06 | 1, 233.64 | 3.05 | 0.08^#^ |
| ^1^Julian Date is scaled from 0 (end of May) to 1 (end of November) | | | | |

| **Table S5:** All corticosterone values, replacing anthelminthic treatment with the number of anthelminthic treatments received prior to each sample, and corticosterone values adjusted for inter-plate variation (n = 268 samples from 100 individuals). | | | | |
| --- | --- | --- | --- | --- |
| **Fixed Effects** | **Estimate ± SE** | **df** | **F-value** | **p-value** |
| Intercept | 8.50 ± 0.58 |  |  |  |
| Food Availability | -0.04 ± 0.23 | 1, 101.18 | 2.86 | 0.09^#^ |
| Egg Burden | -0.09 ± 0.05 | 1, 239.79 | 1.07 | 0.30 |
| Number of Treatments Received | 0.004 ± 0.06 | 1, 103.34 | 0.01 | 0.94 |
| Julian Date^1^ | -2.96 ± 0.59 | 1, 171.43 | 24.68 | <0.0001* |
| Reproductive Status | 0.31 ± 0.17 | 1, 222.10 | 3.35 | 0.07^#^ |
| Sex | 0.57 ± 0.17 | 1, 83.80 | 11.52 | 0.001* |
| Age | 0.79 ± 0.25 | 1, 210.93 | 9.60 | 0.002* |
| Site | 0.30 ± 0.16 | 1, 73.24 | 3.74 | 0.057^#^ |
| Food x Worm Burden | 0.10 ± 0.06 | 1, 234.67 | 2.55 | 0.11 |
| ^1^Julian Date is scaled from 0 (end of May) to 1 (end of November) | | | | |

| **Table S6:** **Linear mixed-effects model of the effect of food supplementation and anthelminthic treatment on wood mouse residual index.** Data is based on 580 samples from 291 individuals, and mouse ID was included as a random effect. Test statistics are calculated using Type II sum of squares and reported with Kenward-Roger degrees of freedom. Reference levels include: no food supplementation, no anthelminthic treatment, and non-reproductive, male, and non-adult individuals. ^#^p < 0.1, *p < 0.05. | | | | |
| --- | --- | --- | --- | --- |
| **Fixed Effects** | **Estimate ± SE** | **df** | **F-value** | **p-value** |
| Intercept | -1.67 ± 0.85 |  |  |  |
| Food Availability | 0.06 ± 0.38 | 1, 332.29 | 0.05 | 0.82 |
| Egg Burden | 0.17 ± 0.08 | 1, 568.94 | 7.05 | 0.008* |
| Drug Treatment | 0.38 ± 0.27 | 1, 248.69 | 1.92 | 0.17 |
| Julian Date^1^ | -1.07 ± 0.86 | 1, 553.79 | 1.55 | 0.21 |
| Reproductive Status | 1.78 ± 0.30 | 1, 569.27 | 35.78 | <0.0001* |
| Sex | 1.20 ± 0.28 | 1, 264.05 | 17.64 | <0.0001* |
| Age | 1.17 ± 0.31 | 1, 555.93 | 13.85 | 0.0002* |
| Site | -0.61 ± 0.27 | 1, 240.71 | 4.97 | 0.03* |
| Food x Worm Burden | -0.46 ± 0.10 | 1, 568.77 | 0.19 | 0.66 |
| ^1^Julian Date is scaled from 0 (end of May) to 1 (end of November) | | | | |

| **Table S7:** **Linear mixed-effects model of the effect of food supplementation and anthelminthic treatment on wood mouse skeletal muscle index.** Data is based on 580 samples from 291 individuals, and mouse ID was included as a random effect. Test statistics are calculated using Type II sum of squares and reported with Kenward-Roger degrees of freedom. Reference levels include: no food supplementation, no anthelminthic treatment, and non-reproductive, male, and non-adult individuals. ^#^p < 0.1, *p < 0.05. | | | | |
| --- | --- | --- | --- | --- |
| **Fixed Effects** | **Estimate ± SE** | **df** | **F-value** | **p-value** |
| Intercept | 16.65 ± 0.96 |  |  |  |
| Food Availability | -0.16 ± 0.43 | 1, 326.92 | 0.29 | 0.59 |
| Egg Burden | 0.13 ± 0.09 | 1, 569.60 | 4.64 | 0.03* |
| Drug Treatment | 0.36 ± 0.31 | 1, 246.17 | 1.38 | 0.24 |
| Julian Date^1^ | 0.79 ± 0.98 | 1, 551.83 | 0.66 | 0.42 |
| Reproductive Status | 1.12 ± 0.34 | 1, 568.12 | 11.03 | 0.001* |
| Sex | 1.42 ± 0.32 | 1, 260.73 | 19.32 | <0.0001* |
| Age | 0.49 ± 0.35 | 1, 553.53 | 1.91 | 0.17 |
| Site | -0.58 ± 0.31 | 1, 237.64 | 3.47 | 0.06^#^ |
| Food x Worm Burden | 0.0001 ± 0.12 | 1, 569.47 | 0.00 | 0.99 |
| ^1^Julian Date is scaled from 0 (end of May) to 1 (end of November) | | | | |
